# Jelly belly: Recovery of fish eDNA from *Cassiopea* medusae gastrovascular cavities across the Florida Keys

**DOI:** 10.64898/2026.08.06.743372

**Authors:** Kaden Muffett, Megan Sporre, Maria Pia Miglietta, Ron Eytan

## Abstract

Ranges of small benthic fauna are notoriously difficult to assess. In some of these cases, modern eDNA methods can shed light on species occurrence. Here we conduct an exploratory study on the fish eDNA recoverable from the gastrovascular cavities of the easy-to-sample pore water siphoning benthic invertebrate, *Cassiopea*, across six sites within the Florida Keys. Twenty-seven fish *12S* identities were recovered from water samples, two from sediment samples, and seventeen from *Cassiopea* gut swabs. In total, thirty-two different species were identified from nineteen families, including one shark species (*Ginglymostoma cirratum*), and five species of cryptobenthic reef fishes (f: Gobiidae, Labrisomidae). Additionally, five species were identified from medusae samples that were not recovered in water or sediment samples. The species identities recovered may provide insight into the fish in direct proximity to *Cassiopea* assemblages, as well as indicate that *Cassiopea* may accrue disproportionate eDNA from cryptobenthic reef fish compared to surrounding environmental samples. The unorthodox sampling technique of using eDNA recovered from jellyfish stomachs yields another avenue for epibenthic community data acquisition.

## Introduction

*Cassiopea*, the Upside-Down Jellyfish, are a common component of mangroves and shallow areas across the Florida Keys and more broadly in the Caribbean and global subtropics. *Cassiopea* engage in mixotrophy, housing algal symbionts while also pulsing water along their bodies and into their oral arms to catch and digest small prey items (Ohdera et al., 2018; Santhanakrishnan et al., 2012).

Like many other gelatinous zooplankters, the dietary profile of *Cassiopea* is poorly described, though previous studies characterizing this in medusae found mainly crustaceans (Larson, 1997; Muffett et al., 2025). Regardless of diet, *Cassiopea* are constantly siphoning water from the benthos around them in a manner that may make their gastrovascular cavities potential reservoirs of DNA and material from the surrounding sediment (Battista et al., 2022; Santhanakrishnan et al., 2012). This may be especially useful in identifying cryptobenthic reef fishes.

Cryptobenthic reef fishes (CRFs) are integral to coastal ecosystems, considered the ‘hidden half’ of reef diversity (Brandl et al., 2018). Identification of these fish in wild environments often relies on ichthyocides and anesthetization stations (Gómez-Buckley et al., 2023). Water sampling for eDNA to identify CRF species has produced results stronger than anesthetic sampling in some cases (Bessey et al., 2023) and low CRF reads relative to destructive sampling methods in others (Gómez-Buckley et al., 2023). Swabbing filter feeders pulling water from the surrounding environs may be a low-cost, low-effort approach to supplement traditional water sampling or time-intensive snorkel and dive searches.

## Methods

### Collection and extraction

*Cassiopea* gastrovascular swabs were collected in August 2021 across eight sites within the Florida Keys, along with 1L water samples (0.2um pore size filter following USGS eDNA sampling protocol) and ∼10 g of surface sediment (see Muffett et al., 2025 for details, see Muffett and Miglietta, 2023 for *Cassiopea* species identification; Fig 1). Samples were stored in 2mL (swab) or 15mL (water filter or sediment sample) dimethyl sulfoxide ethylenediamine tetraacetic acid saturated salt storage solution (1 L pH 7.5: 93.06 g EDTA, 60 mL 20 % NaOH solution, 20 mL 25 % HCl, 40 mL DMSO, 800 mL water, NaCl to oversaturation; see ref. Pavlovska et al., 2021) (commonly referred to as DESS). All sites were shallower than 1.5 m depth.

**Fig 1.**
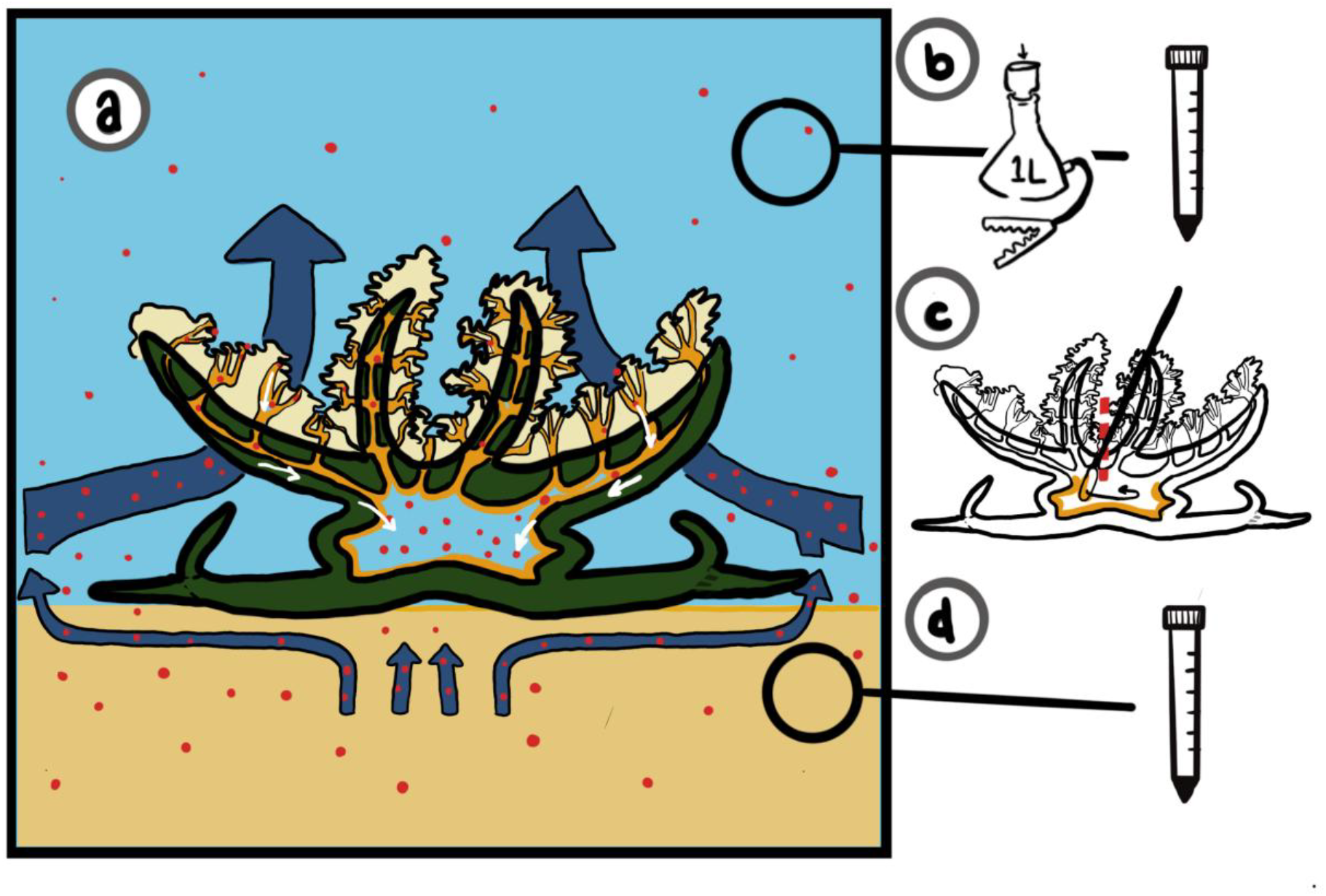
a) Hypothetical feeding mechanism of *Cassiopea* displaying particulate intake. b) Water sampling approach following USGS manual water sampler design with storage in DESS. c) Gastric sampling design, with red dotted line indicating incision point. d) Sediment sampling design (surface sediment sampled directly into DESS).

Samples were retained at room temperature for between 3 and 10 days, then frozen before processing. All samples were then extracted using ZymoBIOMICS DNA Kits (PN# D4300T) following manufacturer instructions.

### PCR and sequencing

PCR reactions were performed using MiFish primers (Miya et al. 2015) and KAPA HiFi HotStart in 8 uL reactions with 4 replicate reactions per sample. PCR products were pooled by sample and a 2 uL aliquot was run on TapeStation. As nonspecific amplification is a known phenomenon with MiFish primers, only samples with amplification in the region of 300 bp exceeding 2 ng/uL were included for purification, then secondary amplification with barcoded secondary primers (following Miya et al., 2015). Two sites failed to amplify and were excluded from sequencing.

After secondary primer appending amplification, samples were sent to Texas A&M AgriLife for sequencing (MiSeq 2×250).

### Data analysis

Raw sequences were uploaded to the MiFish v4.0 online pipeline (Sato et al., 2018) and non-target vertebrates (e.g. human, iguana, pig) were removed from the dataset. To prevent primer-hopping sequences from being included in final sample tallies, any IDs with 230 or below sequences in a given sample (10x the highest number found in the blank) were removed from the dataset. Low-confidence taxonomic identities were confirmed using NCBI BLAST.

## Results

### Dataset

After cleaning, the dataset had an average of 414,917 sequences per sample (min: 56,837, max: 716,180). Out of the 32 medusae, six water samples and six sediment samples checked for presence of 300 bp band, four water samples, one sediment sample and 14 medusa samples were positive for *12S* bands. Across these samples, a total of 32 fish species were identified, with 17 found in gastrovascular cavities at least once (Fig 2). This included 6 species found only in trace abundances (230-1000 reads/sample).

**Fig. 2.**
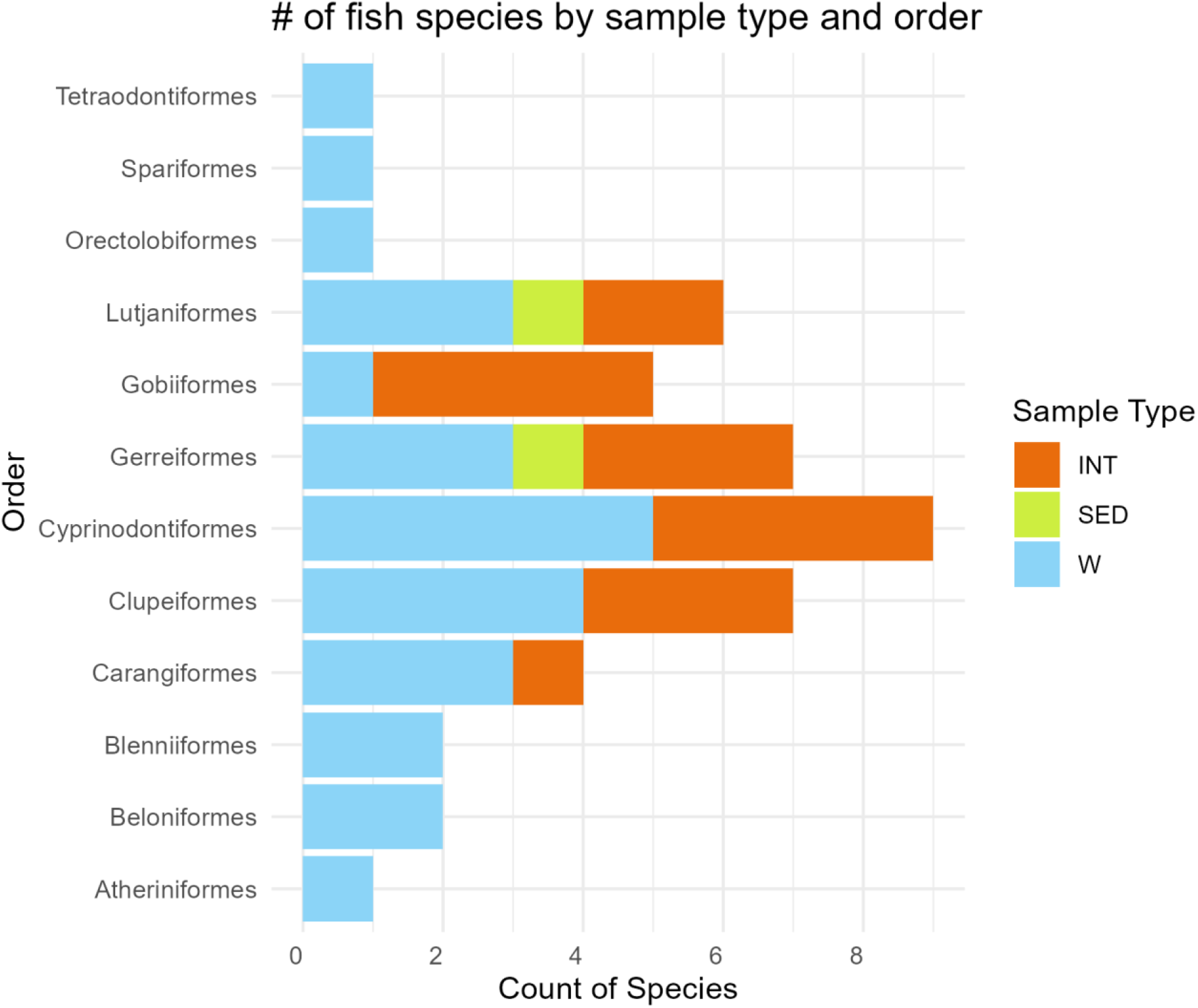
Chart of all fish signatures identified by order and colored by sample type: medusae (orange), water (light blue) and sediment (green).

### CRFs

In successfully amplified water samples, average proportion of reads associated with a member of a canonical CRF family per Brandl 2018 was 1.12% (min: <0.01%, max: 4.34%) with an average number of taxa of 10.5/sample (min: 1, max: 24). In successfully amplified medusa gastrovascular cavities, an average of 40% of reads were from CRF fish (min: <0.01%, max: >99.9%), with an average of 2.14 fish taxa/sample (min: 1, max: 5); however, the number of reads attributable to CRFs was not significantly higher in GVCs than water (Kruskal-Wallis chi-squared = 1.2363, df = 1, p-value = 0.2662). The single successfully amplified sediment sample contained no CRF reads and two total taxa (Fig 3).

**Fig 3.**
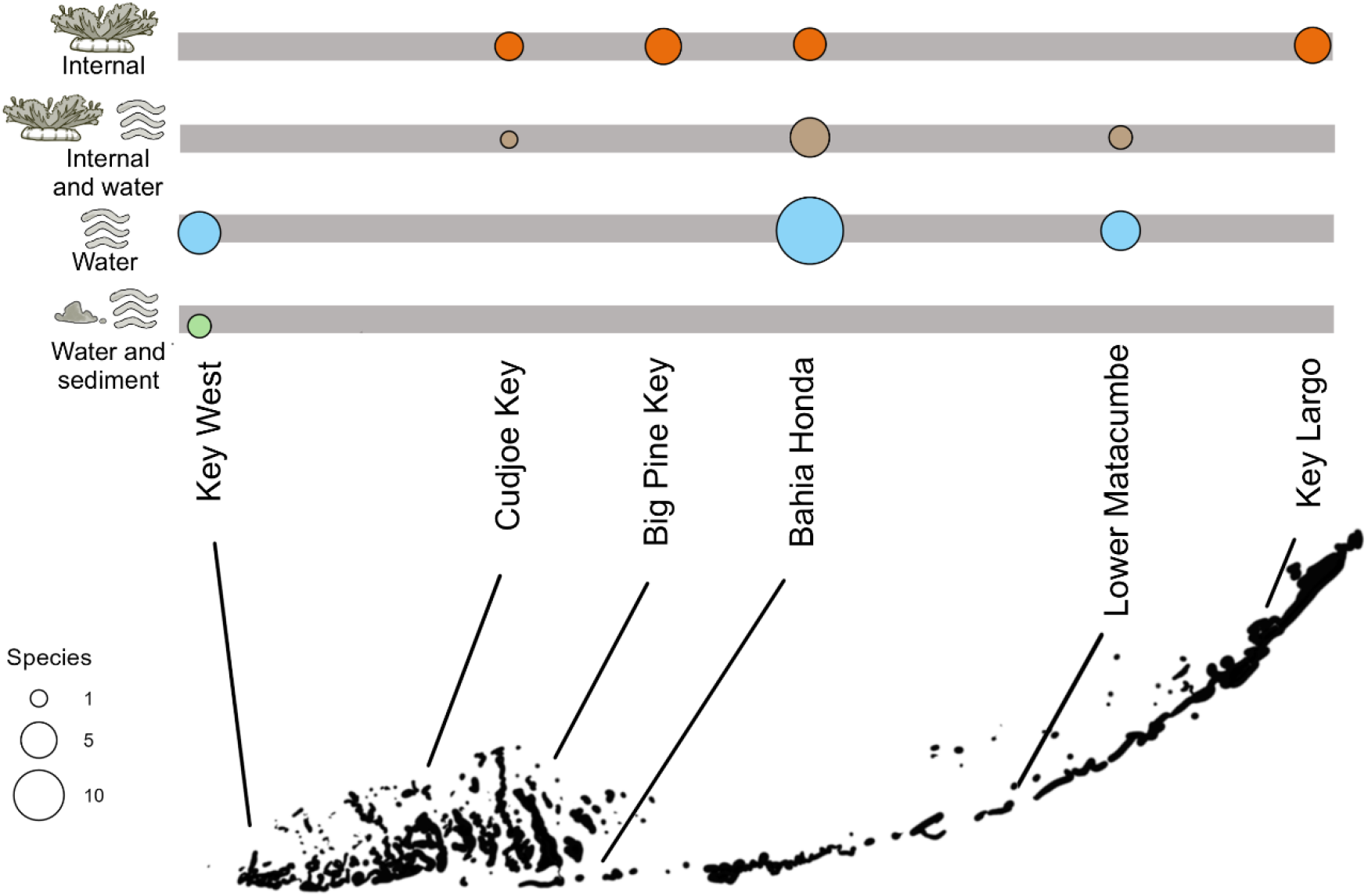
Number of fish species detected by site and sample type. Fish species are separated based on whether they have been found in one (e.g. “Internal”/”Water”) or multiple (e.g. “Internal and water”) sample types. No samples were found in “Sediment” alone or all three sample types together.

Medusae from five sites amplified (all but Key West), but there were no size trends differentiating successfully (7.82 cm diameter ave, 95% CI: 0.16 to 15.5 cm) and unsuccessfully (8.43 cm diameter ave, 95% CI: 1.70 to 15.2 cm) amplifying medusae.

The greatest diversity of eDNA components was found on Bahia Honda Key, with 24 total detections in the water sample and 10 detections within medusa swabs (Fig 4). Four taxa were found in medusa gastrovascular cavities at this site that were not identified in the associated water samples: *Lophogobius cyprinoides, Ctenogobius boleosoma, Caranx crysos*, and *Bathygobius soporator. Lophogobius cyprinoides* was identified in water samples at only one site (Lower Matacumbe Key) but identified in three additional sites using medusae (Big Pine Key, Bahia Honda Key and Key Largo). Water samples and medusae matched imperfectly at all sites, with medusae from two sites capturing taxa lost in water samples, two sites where medusae amplified but water samples did not and three sites where water samples picked up taxa not seen in medusa gastrovascular cavities.

**Fig 4.**
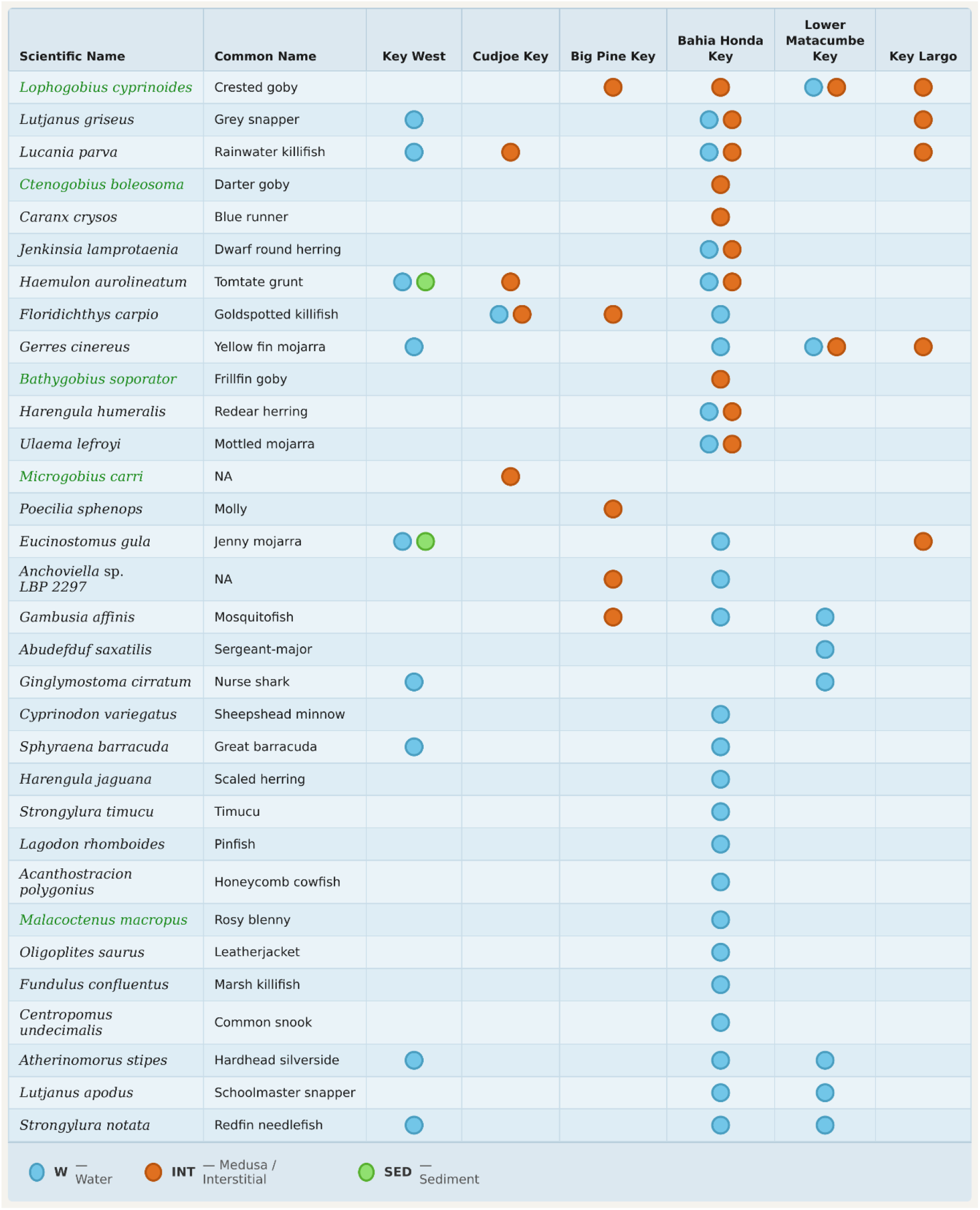
All identified sequences with scientific names, common names and sample types in which fish were detected. Light blue = water samples, orange = medusa samples and green = sediment samples. Cryptobenthic fish are highlighted.

## Discussion

While the described use of cnidarian-derived eDNA is unlikely to become a commonplace sampling technique for fish eDNA work, it brings to light several interesting points. Firstly, eDNA that may be in low abundance in the water column may be in higher abundance within benthic filter feeders. Secondly, two of the species found in medusae but not the surrounding water have been identified as highly abundant in larval fish populations, suggesting that *Cassiopea* may supplement a primarily crustacean diet with larval fish (Gomes et al., 2014). Thirdly, amplifications from gastrovascular cavities were more successful than from associated sediments, even though eDNA residence times within a digestive cavity are likely quite short. While substantial work has demonstrated that fish diets are highly specific, many invertebrates are neglected in complex food web studies (Casey et al., 2019; Hetherington, Choy, et al., 2022; Hetherington, Damian-Serrano, et al., 2022).

The fish recovered within this study are plentiful or well-known residents of the Florida Keys, including schooling fish and benthic fish alike (Ault et al., 2006; Briggs, 1958). For the Grey Snapper (*Lutjanus griseus*) and Jenny Mojarra (*Eucinostomus gula*), juveniles are known to inhabit shallow seagrass beds (Acosta et al., 2007). For the cryptobenthic fish, prevalence within the *Cassiopea* gastrovascular cavity may also be linked to their disproportionate larval supply (Brandl et al., 2019). For sheltering species, for instance, the Rainwater Killifish (*Lucania parva*), local seagrass beds may provide shelter for both the fish and the jellyfish, in the latter case from tidal forces, likely not predation (Jordan, 2002).

The sites of collection in which fish eDNA was only present in *Cassiopea* samples (Big Pine Key and Key Largo) were exceptionally shallow (<20 cm depth at point of collection), while the location with the most water reads (Bahia Honda Key) is a deeper water location ∼1-1.5 m adjacent to both mangroves and a deep-water pass. These medusae were also the largest found outside of enclosed pools. It is possible that the observed pattern of *Cassiopea* reads in locations with low water eDNA is representative of low biomass or high irradiation in these environments (Barnes et al., 2014). As *Cassiopea* themselves engage in active siphoning from the environment for their nutrition and have photoprotective pigments, they serve as reservoirs of animal DNA in these conditions harsh to eDNA (Ohdera et al., 2018).

In this work, we have provided a snapshot into the eDNA found within *Cassiopea* gastrovascular cavities; however, the relationship between medusae and the fish whose eDNA signatures are identified has not yet been established in the literature. While we posit that a likely explanation would be *Cassiopea*’s passive predation on CRF fish eggs and larval forms, with only circumstantial data we cannot rule out that medusae simply accrete local eDNA. *Cassiopea* jellyfish, like other medusae or anemones, may provide refuge in otherwise low-coverage areas (Riascos et al., 2018), or they may simply incidentally consume larval CRFs. Observational work on fish behavior in the vicinity of medusae would better close these knowledge gaps.

## Data availability

All data is available upon request. S Tab 1 provides full species identities by sample.

